# *GATA1* knockout in human pluripotent stem cells generates enhanced neutrophils to investigate extracellular trap formation

**DOI:** 10.1101/2023.02.08.526339

**Authors:** Thomas C. Harper, Elaine M. Oberlick, Tomas J. Smith, Duncan E. Nunes, Mark A. Bray, Seonmi Park, Corey D. Driscoll, Sarah F. Mowbray, Christophe Antczak

## Abstract

Human pluripotent stem cell (hPSC)-derived tissues can be used to model diseases and validate targets in cell types that are challenging to harvest and study at-scale, such as neutrophils. Neutrophil dysregulation, specifically unbalanced neutrophil extracellular trap (NET) formation, plays a critical role in the prognosis and progression of multiple diseases, including COVID-19. hPSCs can provide a limitless supply of neutrophils (iNeutrophils) to study these processes and discover and validate targets *in vitro*. However, current iNeutrophil differentiation protocols are inefficient and generate heterogeneous cultures consisting of different granulocytes and precursors, which can confound the study of neutrophil biology. Here, we describe a method to dramatically improve iNeutrophils’ yield, purity, functionality, and maturity through the deletion of the transcription factor *GATA1. GATA1* knockout (KO) iNeutrophils are nearly identical to primary neutrophils in cell surface marker expression, morphology, and host defense functions. Unlike wild type (WT) iNeutrophils, *GATA1* KO iNeutrophils generate NETs in response to the physiologic stimulant lipopolysaccharide (LPS), suggesting they could be used as a more accurate model when performing small-molecule screens to find NET inhibitors. Furthermore, through CRSPR/Cas9 deletion of *CYBB* we demonstrate that *GATA1* KO iNeutrophils are a powerful tool in quickly and definitively determining involvement of a given protein in NET formation.

## Introduction

Neutrophils are the most abundant immune cells in the human body and make up approximately 70% of circulating leukocytes. They migrate to the site of infections where they recruit other immune cells, and independently destroy invading microorganisms through phagocytosis, the release of granules, and the formation of neutrophil extracellular traps (NETs) (1,2).

While neutrophils are an important first line of defense in the innate immune system, their overactivation can have a proportionally negative impact on many diseases and affect numerous organ systems. The dysregulation of neutrophils is correlated with the progression of multiple diseases including rheumatoid arthritis, atherosclerosis, psoriasis, chronic obstructive pulmonary disease, and gallstone formation (3–7). COVID-19 clearly establishes the link between overactive neutrophils and disease severity and highlights the need for improved methods to model neutrophil dysregulation. SARS-CoV-2, the virus which causes COVID-19, has been shown to directly induce NETs (8,9) and the overproduction of NETs and neutrophil reactive oxygen species (ROS) exacerbate COVID-19 complications including blood clots, cytokine storm, organ damage, and respiratory failure (10–13). Unsurprisingly, there is a strong positive correlation between NET production, disease severity and patient outcome (14,15).

While inhibiting overactive neutrophils has the potential to mitigate COVID-19 severity (15–17), the nature of primary neutrophils severely restricts their utility in drug discovery. Primary neutrophils, like all donor tissues, are limited by access and cross-patient variability, survive *ex vivo* for less than 24 hours, are transcriptionally silent and non-proliferative, and cannot be cryopreserved (18). These shortcomings preclude large-scale drug screening and make unbiased genetic screens and target validation experiments using CRISPR/Cas9 challenging.

Human pluripotent stem cells (hPSCs) provide an inexhaustible source of material that can overcome these challenges. hPSCs self-renew indefinitely, can be differentiated into a variety of highly relevant cell types, and are easy to genetically modify. hPSC-derived cells have been successfully used in a variety of pharmaceutical efforts, ranging from high-throughput phenotypic drug screens to model and correct neurological disorders, to the generation of hepatocytes to screen drug-mediated toxicity (19,20). The production of homogeneous cultures of mature cells is crucial for assay relevance and reproducibility. Unfortunately, current protocols to produce hPSC-derived neutrophils (iNeutrophils) generate a heterogeneous combination of hematopoietic progenitors and cells resembling neutrophils, eosinophils, and basophils (21–25).

Neutrophil specification is tightly regulated by an interplay of cell fate-determining transcription regulators. While eosinophils and neutrophils rely on the expression of *CEBPE* and *GFI1*, post-translational acetylation of CEBPE at K121 and K127 along with the reduction of *GATA1* ultimately determines neutrophil commitment (26,27). During neutrophil maturation, *CEBPE* is downregulated and expression of the terminal granulopoiesis genes *CEBPD* and *SPI1* escalate (28). The small-molecules and cytokines governing these events are largely unknown. Based on transcriptional analysis of sorted iNeutrophils along with mouse genetic studies, we surmised that the deletion of *GATA1*, a transcription factor important for the development of eosinophils and basophils, would force hPSC-derived granulocytes into a neutrophil-specific program and eliminate contaminating cells (29,30). We demonstrate that knocking out *GATA1* in H1 human embryonic stem cells (hESCs) using CRISPR/Cas9 (*GATA1* KO) followed by granulocyte differentiation produces pure populations of iNeutrophils that are nearly identical to their primary counterparts. Compared to wild type (WT) iNeutrophils, *GATA1* KO iNeutrophils have dramatically improved levels of the neutrophil surface markers CD182, CD11b, CD15, CD16 and CD66b and retain their host defense functions including phagocytosis, ROS production and myeloperoxidase (MPO) activity. Unlike WT iNeutrophils, *GATA1* KO iNeutrophils form NETs after treatment with the physiologic NET stimulant lipopolysaccharide (LPS). Furthermore, *GATA1* KO iNeutrophils can be further genetically manipulated through CRISPR/Cas9 to evaluate the role of individual genes in neutrophil functions. *GATA1* KO iNeutrophils with deletion of *CYBB*, which encodes a protein involved in NET formation, produce reduced NETs in response to the NET stimulant phorbol myristate acetate (PMA).

## Materials and Methods

### Neutrophil differentiation

Differentiation was adapted from previously published protocols (31,32). All cells were cultured in humidified tissue culture incubators at 37°C with 5% C0_2_. H1 hESCs were maintained on Matrigel (Corning, 354277) coated dishes using mTeSR Plus media (StemCell Tech, 100-0276). The day before induction (Day -1), cells were dissociated into 100 to 300 μm clumps using ReLeSR (StemCell Tech, 05872) and plated onto hematopoietic Matrigel (Corning, 354234) coated 6 cm dishes in mTeSR Plus media with 10 μM Y-27632 (Tocris, 1254). On Day 0, mTeSR Plus was removed and cells were induced with Base Media 1 (StemPro 34 SFM Media (Gibco, 10639011) containing 1x GlutaMAX (Gibco, 35050079), 0.45 mM aMTG (Sigma-Aldrich, M6145), 200 μg/ml Transferrin (Sigma-Aldrich, 10652202001), and 88 μg/ml L-Ascorbic Acid (Sigma-Aldrich, A4403)) supplemented with 5 ng/ml BMP4 (R&D Systems, 314-BP), 50 ng/ml VEGF (R&D Systems, 293-VE) and 2 μM CHIR99021 (Tocris, 4423). Cells were incubated for two days. On Day 2, cells were refreshed with Base Media 1 plus 5 ng/ml BMP4, 50 ng/ml VEGF, and 20 ng/ml bFGF (R&D Systems, 233-FB). On Day 4, cells were again refreshed with Base Media 1 plus 15 ng/ml VEGF and 5 ng/ml bFGF. On Day 6, media was replaced with Base Media 2 (StemPro 34 SFM Media containing 1x GlutaMAX, 0.4 mM aMTG, 150 μg/ml Transferrin, and 50 μg/ml L-Ascorbic Acid)) supplemented with Growth Factor Cocktail (5 ng/ml BMP4, 5 ng/ml bFGF, 10 ng/ml IL-6 (R&D systems, 206-IL), 10 ng/ml Flt3L (PeproTech, 300-19), 5 ng/ml IL-11 (PeproTech, 200-11), 25 ng/ml IGF-1 (R&D Systems, 291-G1), 30 ng/ml IL-3 (R&D Systems, 203-IL), 30 ng/ml TPO (PeproTech, 300-18), 50 ng/ml VEGF, 100 ng/ml SCF (R&D Systems, 255-SC), 20 ng/ml SHH (PeproTech, 100-45) and 10 μg/ml Angiotensin II (Sigma-Aldrich, A9525)). After one or two days, cells began to shed away from the adherent monolayer, so during each media change, used media containing floating cells was collected, centrifuged at 300 x g for 10 minutes to pellet cells, resuspended in fresh media, and returned to their original dishes. Media was changed every other day using Base Media 2 plus Growth Factor Cocktail until Day 12, when floating cells were collected and plated into new 6 cm dishes with Maturation Media (RPMI (Gibco, 61870036), 10% FBS, 1 mM Sodium Pyruvate (Gibco, 11360070), 55 μM b-Mercapto, 25 mM HEPES (Gibco, 15630130), and 100 ng/ml G-CSF (Peprotech, 300-23)). Maturation Media was changed every other day until Day 18 when G-CSF was removed for 24 hours before experiments on Day 19 to avoid G-CSF related stimulation. At harvest, cells were strained through a 15 μm filter (pluriSelect, 43-50015-03) to remove debris followed by centrifugation for 10 minutes at 300 x g to pellet cells. Cell numbers were quantified using a Beckman Coulter Vi-Cell XR cell viability analyzer.

### *GATA1* KO using CRISPR/Cas9

An Amaxa Nucleofector II Device and Nucleofector Kit (Lonza, VPH-5012) were used to transiently express 5 μg of *GATA1* gRNA plasmid (gRNA sequence: ggtgtggaggacaccagagcagg) containing a puromycin resistance gene into 1 × 10^6^ H1 hESCs constitutively expressing *Cas9* from the *AAVS1* locus. After nucleofection, cells were plated onto a 6 cm dish in mTeSR plus media with 10 μM Y-27632, and two days later selected using 1 μg/ml puromycin for 2 days, clonally expanded, genomic DNA extracted (Invitrogen, K1820-1), and target locus PCR amplified (forward primer: gatgcaggagggaaaagagagga, reverse primer: gcaaccaccacatacttccagt) using Platinum *Taq* DNA Polymerase (Invitrogen, 11304011). Amplicons were analyzed using Sanger sequencing and clones with frame shift deletions picked for expansion. All experiments were performed with a *GATA1* KO clone containing a 13 base-pair frame shift deletion (Figure S1). *GATA1* is located on the X-chromosome so only one allele of H1 hESCs (XY karyotype) required editing.

### Hematoxylin and eosin staining

To visualize morphology, 1.5 × 10^5^ cells were suspended in 200 μl PBS plus 1% BSA (Sigma-Aldrich, A9576) and spun onto glass slides using a Thermo Scientific Cytospin 4 centrifuge at 300 x g for 5 minutes, processed through a Siemens Hematek 2000 for staining and sealed / preserved using DPX mounting media (Sigma-Aldrich, 06522). Stained cells were then visualized using a Nikon Eclipse Ci-L microscope.

### Flow cytometry

5 × 10^5^ cells per condition were distributed into a 96-well v-bottom plate (Corning, 3357), washed with stain buffer (BD Biosciences, 554657), then incubated with 200 μl fluorophore-conjugated antibodies diluted in stain buffer for 30 minutes on ice. Cells were then washed two times and fixed with 4% paraformaldehyde (Electron Microscopy Services, 15710) diluted in PBS for 30 minutes followed by two more washes. For washes, 200 μl of stain buffer were added to each sample well followed by centrifugation at 300 x g for 5 minutes at 4°C. All steps were performed in dark conditions and on ice. Antigen specific antibodies reacted to UltraComp eBeads (Invitrogen, 01-3333-42) were used as single-color compensation controls and corresponding isotype controls were included for each antigen specific antibody. Experiment was run using a Yeti (Propel Labs) flow cytometer and analysis was performed using FlowJo software. To calculate positive marker expression, a cut off of no more than 3% isotype background was used. A list of antibodies used is available in Supplemental Tables S1A and S1B.

### Cell sorting

10 × 10^6^ Day 19 WT iNeutrophils were resuspended in 4 ml PBS + 1% BSA (Sigma-Aldrich, A9576) and sorted using a FACSAria III (BD Biosciences) based on low and high forward- and side-scatter populations gated in Figure 1. Once sorted, the two populations were immediately processed for RT-qPCR.

**Fig. 1.**
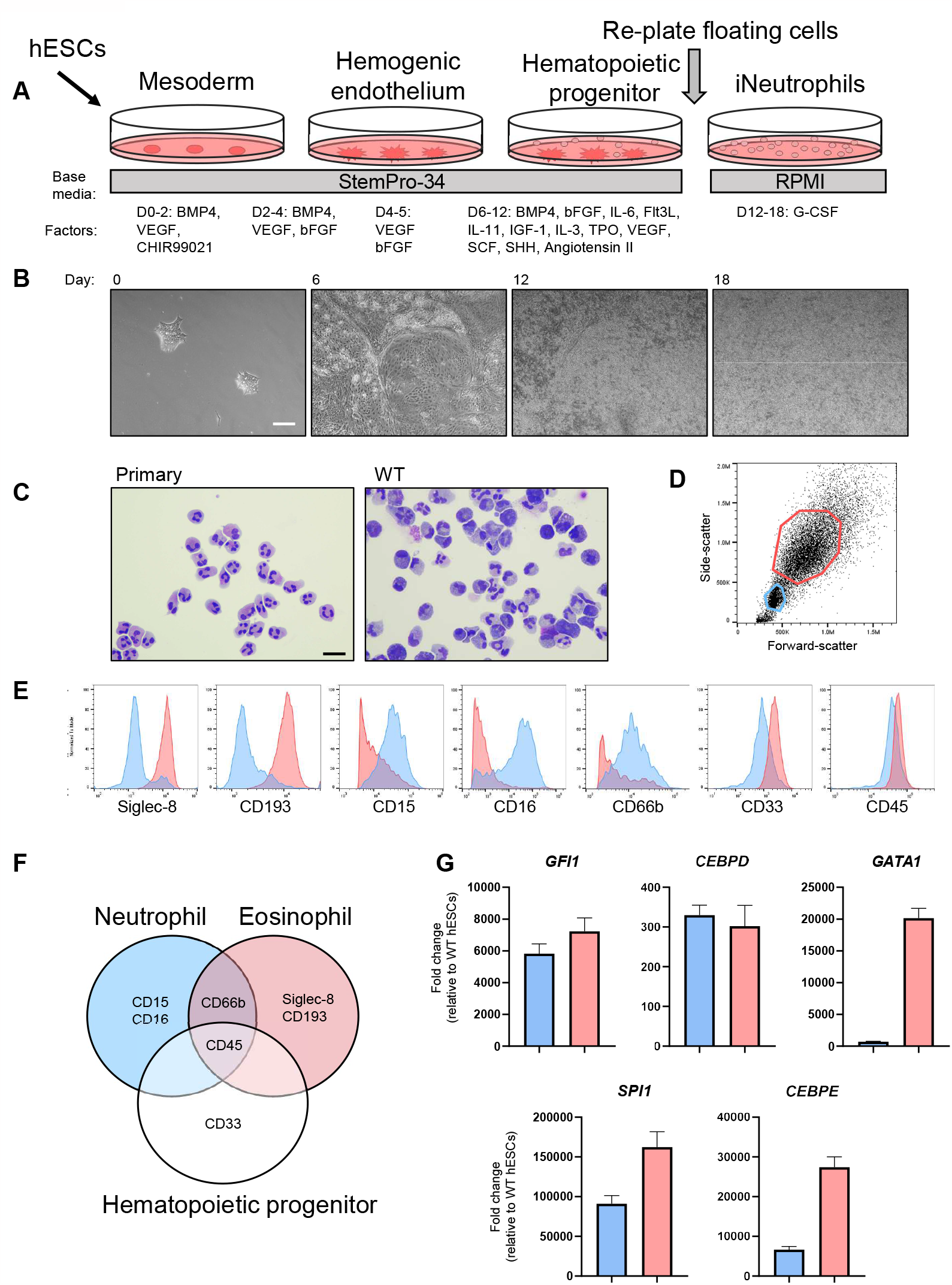
iNeutrophils generated using conventional differentiation methods are heterogeneous. (A) Schematic timeline of differentiation process including base media and soluble factor supplements. (B) Bright field images of differentiation at key timepoints. Scale bar = 100 μm. (C) Hematoxylin and eosin images of iNeutrophils harvested on Day 19 of differentiation compared to primary neutrophils. Scale bar = 20 μm. (D) Flow cytometry scatter plot of iNeutrophils collected on Day 19 of differentiation. Red gate surrounds high forward- and side-scatter population and blue gate surrounds low forward- and side-scatter population. (E) Flow cytometry analysis of blue and red gated populations from scatter plot. (F) Venn-diagram of neutrophil, eosinophil, and hematopoietic progenitor cell surface markers. (G) RT-qPCR of *GFI1, CEBPD, GATA1, SPI1* and *CEBPB* performed on blue and red gated populations from (E) after sorting using FACS. Data normalized to GAPDH (n = 1 independent experiment; mean ± SD of technical triplicates).

### RT-qPCR

Approximately 2 × 10^6^ cells were harvested, and RNA extracted and purified using the RNeasy Mini Kit (Qiagen, 74104), reverse transcribed using SuperScript IV Reverse Transcriptase (Invitrogen, 18090010), and gene expression analyzed by TaqMan assay using Fast Advanced Master Mix (Applied Biosystems, 4444963) and the QuantStudio Flex Real-Time PCR System (Applied Biosystems, 4485701). Fold-changes relative to WT H1 hESCs were calculated using the delta-delta Ct method and normalized using the housekeeping gene *GAPDH* and experimental error was calculated through standard deviation (33). For the time-course study, samples were collected from three independent differentiations and performed in technical triplicates. TaqMan probes used are listed in Supplemental Table S2.

### ROS production

ROS release was measured using the CM-H2DCFDA (General Oxidative Stress Indicator) kit (Invitrogen, C6827) following the manufacturer’s protocol. Briefly, cells were resuspended to 1×10^6^ cells/ml in warm Hanks Balanced Salt Solution (HBSS, Gibco, 14025-092) with 5 μM CM-H2DCFDA dye, incubated at 37°C for 20 minutes, then washed with HBSS. Sotrastaurin was transferred to black-walled, clear-bottomed 384-well cell plates (Greiner, 781091) using the LabCyte Echo for a final concentration of 30 μM, then 2.5 ×10^4^ cells / well were added directly into plates and pre-treated for 30 minutes at 37°C. Plates were washed with HBSS, then treated with 10 nM PMA stimulant or DMSO for one hour at 37°C. Fluorescence of 2’,7’-dichlorofluorescein generated by ROS-induced oxidation of the DCFDA reagent to was measured on a CLARIOstar plate reader at 488 / 535 excitation / emission, then adjusted down by 75% of highest well to bring all wells into range. Mean fluorescence from cell-free wells was subtracted to control for background fluorescence. Experiment was performed on four independent differentiations and three independent donors in at least five technical replicates per experiment.

### Primary neutrophil isolation

Peripheral blood from healthy donors (defined as not having asthma or allergies and not having taken NSAIDS within the previous 5 days) was obtained at Novartis Institutes of Biomedical Research using informed consent under an approved Institutional Review Board research protocol. Fresh blood was EDTA anti-coagulated and used within two hours of donation. Primary donor neutrophils were extracted using Ficoll-density centrifugation. Per 10 ml of fresh blood, 5 ml of PBS and 5 ml of 4% Dextran (Sigma-Aldrich, 31392-50G) in PBS (Gibco, 10010-023) were added and mixed in a 50 ml tube by gently by inverting 2.5 times, then allowed to settle for 30 minutes at room temperature, separating into a dense layer topped with a supernatant containing leukocytes. 75% of the supernatant volume of Ficoll-Paque Premium (Sigma-Aldrich, GE17-5442-03) was added to a new 50 ml falcon tube. The supernatant was carefully transferred on top of the Ficoll, then centrifuged at 650 x g for 20 minutes at room temperature, with a low acceleration (2) and no brake (deceleration set to 0). The supernatant was removed, then the pellet was resuspended in 10 ml of water (Ultrapure diH2O) and mixed no more than 30 seconds to lyse red blood cells. Then 10 ml of 2 x PBS (made from 10 x PBS Gibco, 70011044) was added, and tubes were centrifuged 300 x g for 10 minutes at room temperature (reset acceleration and deceleration to 9).

The supernatant was aspirated, and the pellet containing granulocytes was resuspended in IMDM (Gibco, 21056023) and counted with a ViCell Cell Counter. For the ROS assay, primary neutrophils were resuspended in Hanks Balanced Salt Solution (HBSS, Gibco, 14025-092) and counted.

### Phagocytosis

Primary neutrophils or iNeutrophils were seeded in V-bottom 96-well plates (Greiner, 651201) at 5 × 10^5^ cells / well in 100 μl of IMDM (Gibco, 21056023), then treated with either DMSO or 20 μM cytochalasin D (Sigma-Aldrich, C8273) and incubated at 37°C for 1 hour. During the incubation, 500 μl of Fluoresbrite YG Carboxylate Microspheres 1.75 μm (PolySciences, 17687-5) were opsonized in 4500 μl of Normal Human Serum (GeminiBio, 100-110) diluted 50 / 50 in PBS. Fluorescent particles were incubated at room temperature in the dark for 5 minutes then spun down at 300 x g. The supernatant was removed and the opsonization process was repeated one more time. After the second spin the beads were resuspended in 5 ml of 50 / 50 normal human serum / PBS mixture and 100 μl of the opsonized beads were added per well and incubated for 30 minutes at 37°C. After incubation, the cells were spun down at 300 x g in a 4°C centrifuge and the supernatant was removed. Cells were fixed for 5 minutes with 100 μl / well 4% paraformaldehyde in PBS, then washed twice with ice-cold PBS. After the second wash the supernatant was removed and 200 μl of PBS, followed by 100 μl of 0.4% trypan blue, was added to each well. Cells were then analyzed by flow cytometry for uptake of fluorescent particles. Experiment was performed on at least three independent differentiations and three independent donors in at least technical triplicates.

### MPO activity

The MPO activity of cell lysates was measured using an EnzChek Myeloperoxidase Activity Assay Kit (Invitrogen, E33856) following manufacturer’s instructions. Briefly, cells were resuspended in at 5 × 10^5^ cells / ml in PBS and lysed through freeze thaw cycles and 25 ul added to each well of a 384-well dish (PerkinElmer, 6007270). Chlorination was measured by addition of AFP reagent and fluorescence was measured using a BMG PHERAstar at excitation and emission wavelengths of 485 nm and 520 nm, respectively. Mean fluorescence from cell-free wells was subtracted from experiment wells to control for background fluorescence. Experiments were performed on three independent differentiations and three independent donors using at least technical triplicates.

### NET formation and small molecule inhibition

iNeutrophils or primary neutrophils were plated in IMDM (Gibco, 21056023) at a density of 5 × 10^4^ cells per well in ultra-low base 384-well dishes (Aurora, ABD241001A) and stimulated using DMSO, 50 nM phorbol myristate acetate (PMA, Sigma-Aldrich, P1585), 25 μg/ml Lipopolysaccharide (LPS) O128:B12, (Sigma-Aldrich, L2887) or 5 μM A23187 (Sigma-Aldrich, C7522) for 3 hours at 37°C. For inhibition studies, cells were pre-treated with DMSO, 20 μM sotrastaurin (Selleck Chemicals, S2791), 20 μM diphenyleneiodonium chloride (DPI, Sigma-Aldrich, D2926), 100 μM 4-aminobenzoic acid hydrazide (4-ABAH, Sigma-Aldrich, A41909) or 20 μM disulfiram (Tocris, 3807) for 1 hour followed by 3 hours of stimulation using 50 nM phorbol myristate acetate (PMA, Sigma-Aldrich, P1585), 25 μg/ml Lipopolysaccharide (LPS) O128:B12, (Sigma-Aldrich, L2887) or 5 μM A23187 (Sigma-Aldrich, C7522) at 37°C. After the 3 hour stimulation, a fix/perm/stain solution was added for a final concentration of 2% paraformaldehyde (Electron Microscopy Services, 15710), 0.1% Triton ×100 (Sigma-Aldrich, ×100-100ML) and 50 nM Sytox Green (Invitrogen, S7020). Nine fields per well were imaged using the Yokogawa CV8000 automated microscope at 20x magnification. Image features were extracted using CellProfiler (version 4.2.4) followed by analysis using custom supervised machine learning software to classify NET versus non-NET nuclei based on nuclei features including size, shape, intensity, etc. Experiments were performed on at least three independent differentiations and three independent donors in at least technical triplicates.

## Results

### Conventional cytokine differentiation yields heterogeneous iNeutrophils

First, we generated hemogenic endothelium using cytokines and small molecules following previously published protocols (28,29) (Figures 1A and 1B). Next, we supplemented the hematopoietic progenitor’s media with G-CSF to push the cells towards the neutrophil lineage (34). Hematoxylin and eosin images of Day 19 iNeutrophils revealed a variety of cell types with morphologies consistent with different granulocytes and progenitors (Figure 1C). Flow cytometry analysis identified two major populations distinguishable by size (forward-scatter) and granularity (side-scatter) (Figure 1D). The larger, more granular cells expressed high levels of the non-neutrophil granulocyte surface markers Siglec-8 and CD193, while the smaller, less granular cells expressed high levels of the neutrophil surface markers CD15 and CD16. These smaller, less granular cells also expressed lower levels of the hematopoietic progenitor marker CD33 and higher levels of the mature granulocyte marker CD66b, suggesting this population is immunophenotypically like mature neutrophils. As expected, the pan-hematopoietic marker CD45 was similar in both populations of floating cells (Figures 1E and 1F).

Next, we sorted the low and high forward- and side-scatter populations using FACS and compared the transcript levels of five regulators of granulocyte specification. While most regulators had a five-fold difference or less in transcript levels between the two groups, *GATA1* was upregulated more than 25-fold in the non-neutrophil population (Figure 1G). Studies in mice demonstrate that *Gata1* is critical in the development of eosinophils and basophils, and while it is expressed in the common myeloid progenitor, it is dispensable for the differentiation and function of neutrophils (29,30). These findings suggest that *GATA1* is a key gene responsible for specifying the non-neutrophil population and downregulation could restrict cells toward the desired neutrophil fate.

### *GATA1* KO improves iNeutrophil specification

We devised a novel differentiation approach by deleting *GATA1* in our hESCs to restrict their differentiation capacity to the desired neutrophil cell type. Like their WT counterparts, *GATA1* KO hESCs were able to self-renew and expressed high levels of the pluripotency markers *OCT4* and *NANOG* (Figure 2A). Upon differentiation, both the WT and *GATA1* KO cells downregulated these pluripotency genes and began expressing the hematopoietic transcription regulators *SPI1* and *GFI1* (Figure 2B). By Day 12, the WT and *GATA1* KO hematopoietic progenitors showed differences in gene expression suggesting the *GATA1* KO cells were more neutrophil-like than the WT cells. The Day 12 *GATA1* KO hematopoietic progenitors expressed significantly higher levels of the neutrophil genes *AZU1, AQP9, ELANE* and *MPO*, and significantly lower levels of the eosinophil and basophil-specific gene *CLC* (Figures 2C and 2D). As seen with primary neutrophils, Day 19 WT and *GATA1* KO iNeutrophils expressed low levels of mRNA which prevented us from comparing gene expression with early time-points (35,36).

**Fig. 2.**
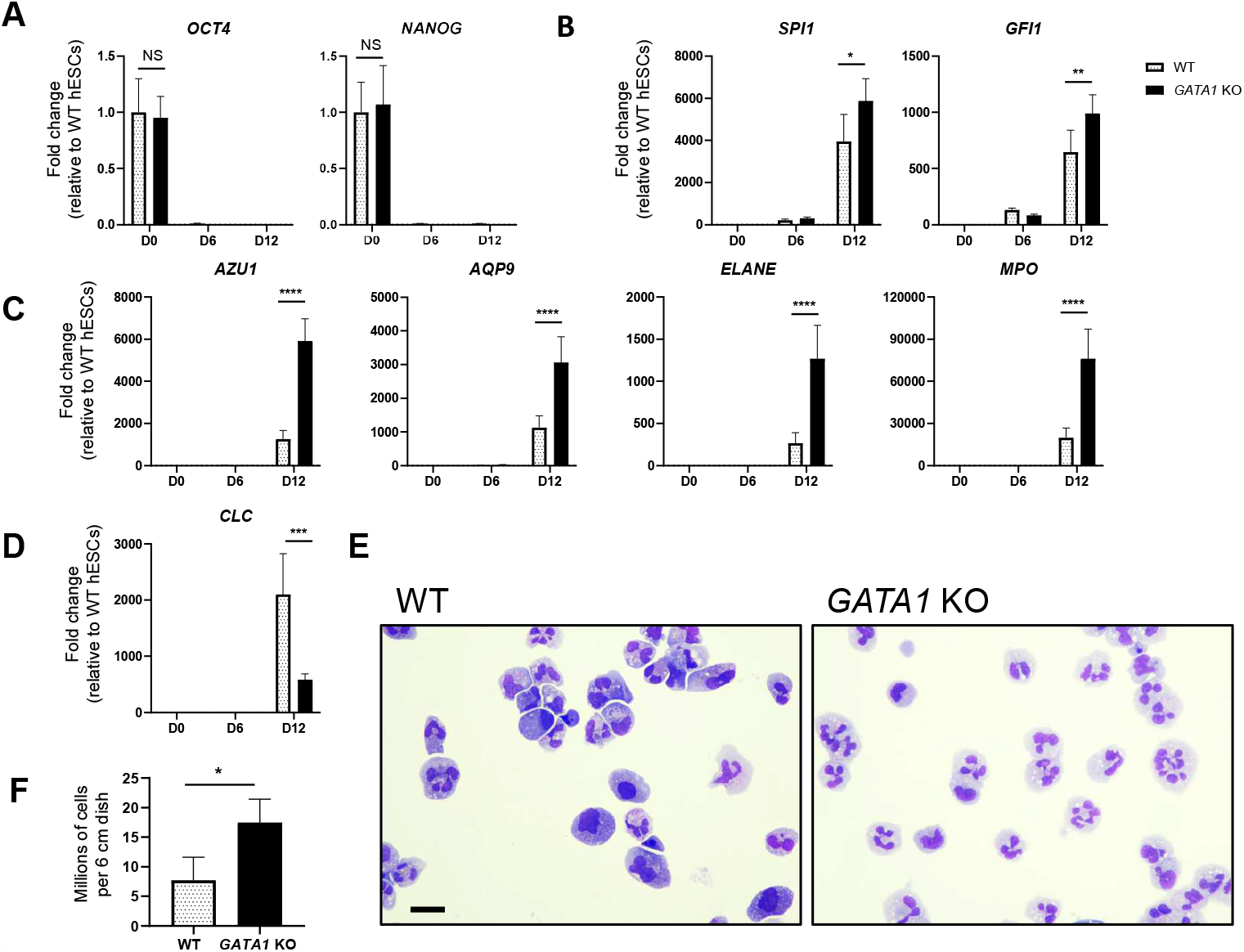
*GATA1 KO* improves iNeutrophil gene expression and phenotype. (A) RT-qPCR of pluripotency genes *OCT4, NANOG*, (B) hematopoietic transcription regulators genes *SPI1, GFI1*, (C) neutrophil specific genes *AZU1, AQP9, ELANE, MPO* (D) and the eosinophil and basophil gene *CLC* on Day 0, 6 and 12 of differentiation. Data is normalized to GAPDH and expressed as fold-change relative to WT hESCs (n = 3 independent experiments; mean ± SD). *P < 0.05, **P < 0.01 ***P < 0.001, ****P < 0.0001 determined by two-way ANOVA. (E) Hematoxylin and eosin images of WT and *GATA1* KO iNeutrophils harvested on Day 19. Scale bar = 20 μm. (F) Number of floating cells produced by WT and *GATA1* KO cultures in each 6 cm dish on Day 19 (n = 4 independent experiments; mean ± SD). *P < 0.05 determined by unpaired t-test.

Hematoxylin and eosin images of Day 19 *GATA1* KO iNeutrophils showed a dramatic increase in the number of cells with the classic neutrophil multilobulated nuclear morphology (Figures 2E and S2).

Furthermore, the *GATA1* KO cells generated on average 17 × 10^6^ cells from each 6 cm dish, more than double that of the WT cells, demonstrating that this improved differentiation method can produce at-scale numbers of homogeneous iNeutrophils (Figure 2F).

### *GATA1* KO iNeutrophils share many characteristics of primary neutrophils

Surface proteins on immune cells mediate cell communication and signal transduction and are often used to distinguish different granulocytes. We used fluorophore-conjugated antibodies specific to basophil, eosinophil, and neutrophil surface proteins (Figure 3A) and flow cytometry to compare WT and *GATA1* KO iNeutrophils versus primary neutrophils. Staining WT iNeutrophils using antibodies against Siglec-8 showed that roughly 50% of the cells adopted an eosinophil phenotype, supported further by the co-expression of CD193 in 25% of the total floating cells (Figures 3B and S3A). Additionally, 26% of the WT cells expressed the non-neutrophil granulocyte marker CD49d. Alternatively, 5% of the *GATA1* KO iNeutrophils expressed Siglec-8 and 4% co-expressed CD193. Only 6% of the *GATA1* KO iNeutrophils expressed CD49d (Figures 3B and S3A).

**Fig. 3.**
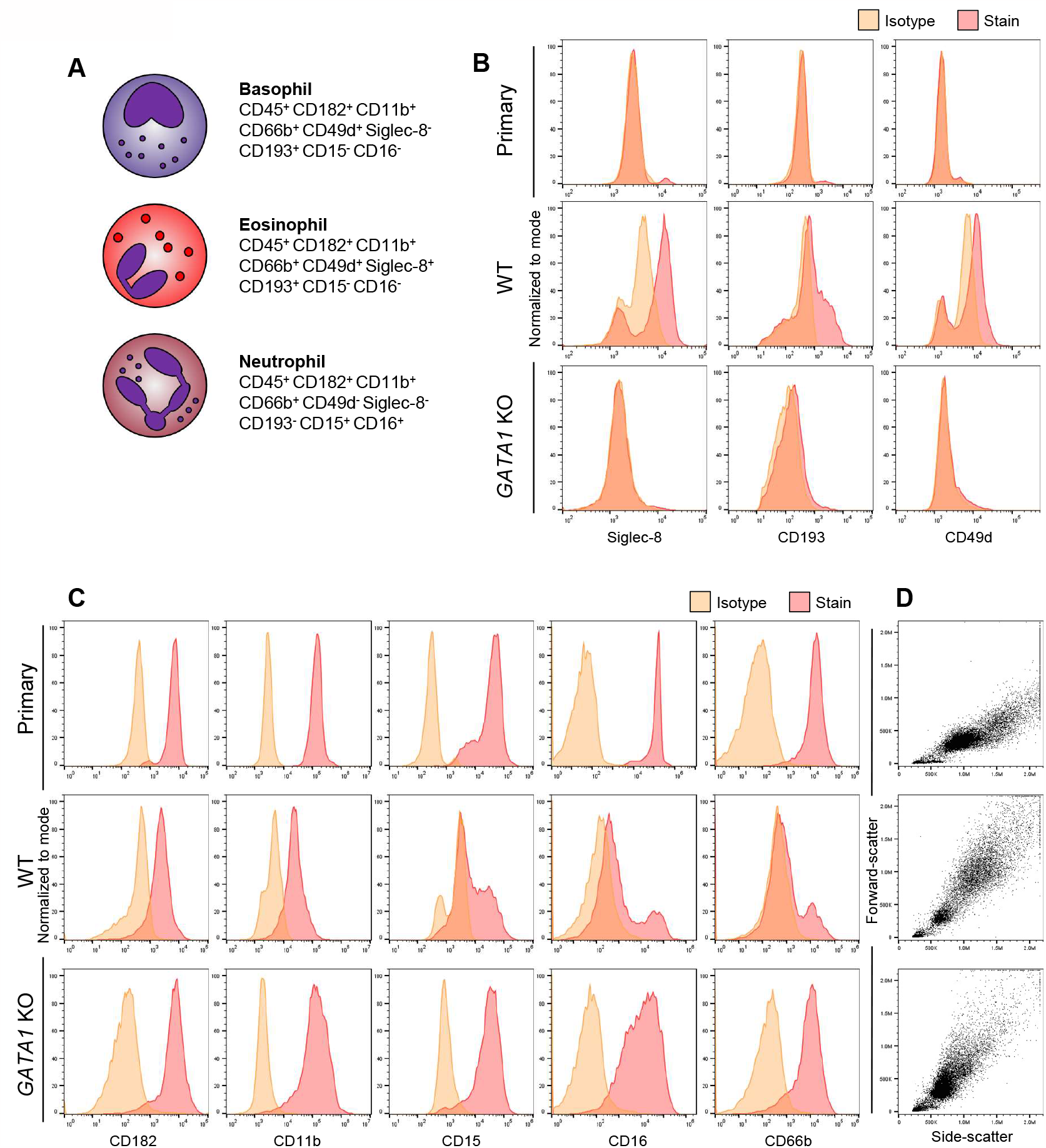
*GATA1* KO iNeutrophils are immunophenotypically like primary neutrophils. (A) Schematic summary of proteins expressed on basophils, eosinophils and neutrophils. (B) Flow cytometry analysis of primary neutrophils, WT and *GATA1* KO iNeutrophils stained using antibodies against non-neutrophil granulocyte surface markers Siglec-8, CD193 and CD49d (C) and antibodies against neutrophil markers CD182, CD11b, CD15, CD16 and CD66b on Day 19. Plots show isotype control (orange) and corresponding specific antibody (red) histograms. (D) Flow cytometry generated forward- and side-scatter plots of primary neutrophils, WT and *GATA1* KO iNeutrophils on Day 19.

While more than 80% of the WT iNeutrophils are CD182 and CD11b positive, fewer expressed the mature neutrophil markers CD15 (50%) and CD16 (29%), and only 22% express the mature granulocyte marker CD66b (Figure 3C).

Alternatively, we saw a dramatic increase in not only the number of CD182 (94%) and CD11b (98%) positive *GATA1* KO iNeutrophils, but also in the magnitude of signal (Figure 3C). More than 90% of the *GATA1* KO iNeutrophils expressed CD15 and CD16, and more than 85% expressed CD66b. Multiplexed staining revealed that more than 80% of the *GATA1* KO iNeutrophils were co-positive for the neutrophil surface markers tested compared to 14% of the WT iNeutrophils (Figure S3B). A list of surface proteins and expression percentages for primary neutrophils, WT and *GATA1* KO iNeutrophils is available in Supplemental Table S3.

The *GATA1* KO iNeutrophils produced a homogeneous forward- and side-scatter profile which largely localized to the previously determined neutrophil-like cell population seen in WT cells (Figures 1D, 1E, and 3D). The diffuse, non-neutrophil like population was dramatically reduced. Interestingly, the *GATA1* KO iNeutrophil population overlaps with the forward- and side-scatter profile seen in primary neutrophils (Figure 3D). Taken together, these results clearly show a remarkable similarity in surface protein expression, size, and granularity between the *GATA1* KO iNeutrophils and primary neutrophils.

### *GATA1* KO does not impact host defense functions

Neutrophils are a critical component of innate immunity and kill invading microorganisms through phagocytosis, MPO release, and ROS production. Analysis revealed that the *GATA1* KO iNeutrophils retained these important functions.

The WT and *GATA1* KO iNeutrophils were able to phagocytose human serum opsonized fluorescent microspheres *in vitro;* however, uptake in the WT cells was reduced relative to the *GATA1* KO cells (22 ± 8% vs. 41 ± 14%, respectively). While the *GATA1* KO iNeutrophils had a slightly lower rate of phagocytosis relative to the primary neutrophils (46 ± 29%), the *GATA1* KO iNeutrophils were less variable. As expected, baseline phagocytosis was inhibited in all groups after treatment with the actin polymerization inhibitor cytochalasin D (Figures 4A and 4B).

**Fig. 4.**
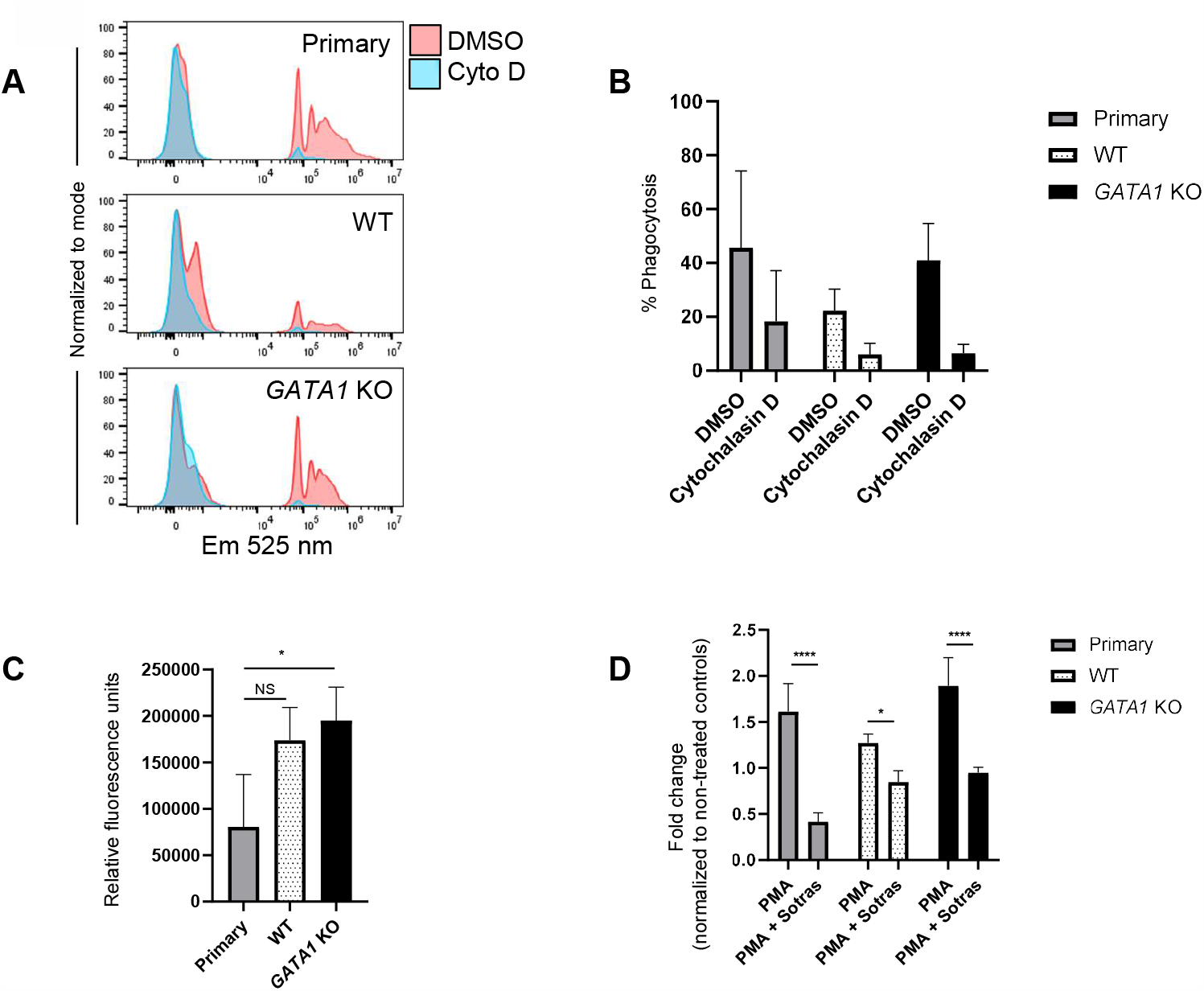
*GATA1* KO iNeutrophils retain critical host defense functions. (A) Histograms represent phagocytosis of opsonized fluorescent microspheres in primary neutrophils, WT and *GATA1* KO iNeutrophils on Day 19 treated with DMSO (red) or 20 μM cytochalasin D (blue) along with (B) percentage uptake (n = 3 independent experiments and 3 independent donors; mean ± SD). (C) Myeloperoxidase (MPO) activity tested via chlorination in primary neutrophils, WT and *GATA1* KO iNeutrophils on Day 19 (n = 3 independent experiments and 3 independent donors; mean ± SD). *P < 0.05 determined by one-way ANOVA. (D) Reactive oxygen species (ROS) release in primary neutrophils, WT and *GATA1* KO iNeutrophils on Day 19 after treatment with DMSO, 10 nM PMA or 10 nM PMA + 30 μM sotrastaurin. Values are presented as fold-change of mean fluorescent intensity relative to DMSO treated cells (n = at least 3 independent experiments and 3 independent donors; mean ± SD). ****P < 0.0001, *P < 0.05 determined by two-way ANOVA.

The *GATA1* KO iNeutrophils retained their MPO activity to the same degree as WT iNeutrophils, but at elevated levels relative to primary neutrophils (Figure 4C). Additionally, WT and *GATA1* KO iNeutrophils along with primary neutrophils generated baseline ROS and were further stimulated through treatment with 10 nM phorbol myristate acetate (PMA). As expected, ROS production was inhibited with the selective protein kinase C (PKC) inhibitor sotrastaurin in all groups (Figure 4D).

### *GATA1* KO iNeutrophils form NETs like primary neutrophils

The formation of NETs was assessed in the WT and *GATA1* KO iNeutrophils after stimulation with PMA, the calcium ionophore A23187, and the bacterial toxin LPS. These stimulants were chosen because they induce NETs using diverse pathways. Treatment with 50 nM PMA stimulated NETs in similar numbers of WT (50 ± 5%), *GATA1* KO iNeutrophils (51 ± 13%), and primary neutrophils (67 ± 4%). Similarly, A23187 was able to induce NETs in both WT and *GATA1* KO iNeutrophils, while WT iNeutrophils produced more NETs after stimulation (83 ± 4%) compared to both *GATA1* KO iNeutrophils (62 ± 6%) and primary neutrophils (51 ± 8%) (Figures 5A and S4A). Considering both PMA and A23187 also generate extracellular traps (ETs) in other granulocytes, and flow cytometry determined that roughly 50% of the WT cells expressed the eosinophil surface marker Siglec-8, the large number of ETs seen in the WT iNeutrophils could be a product of non-neutrophil stimulation (37,38).

**Fig. 5.**
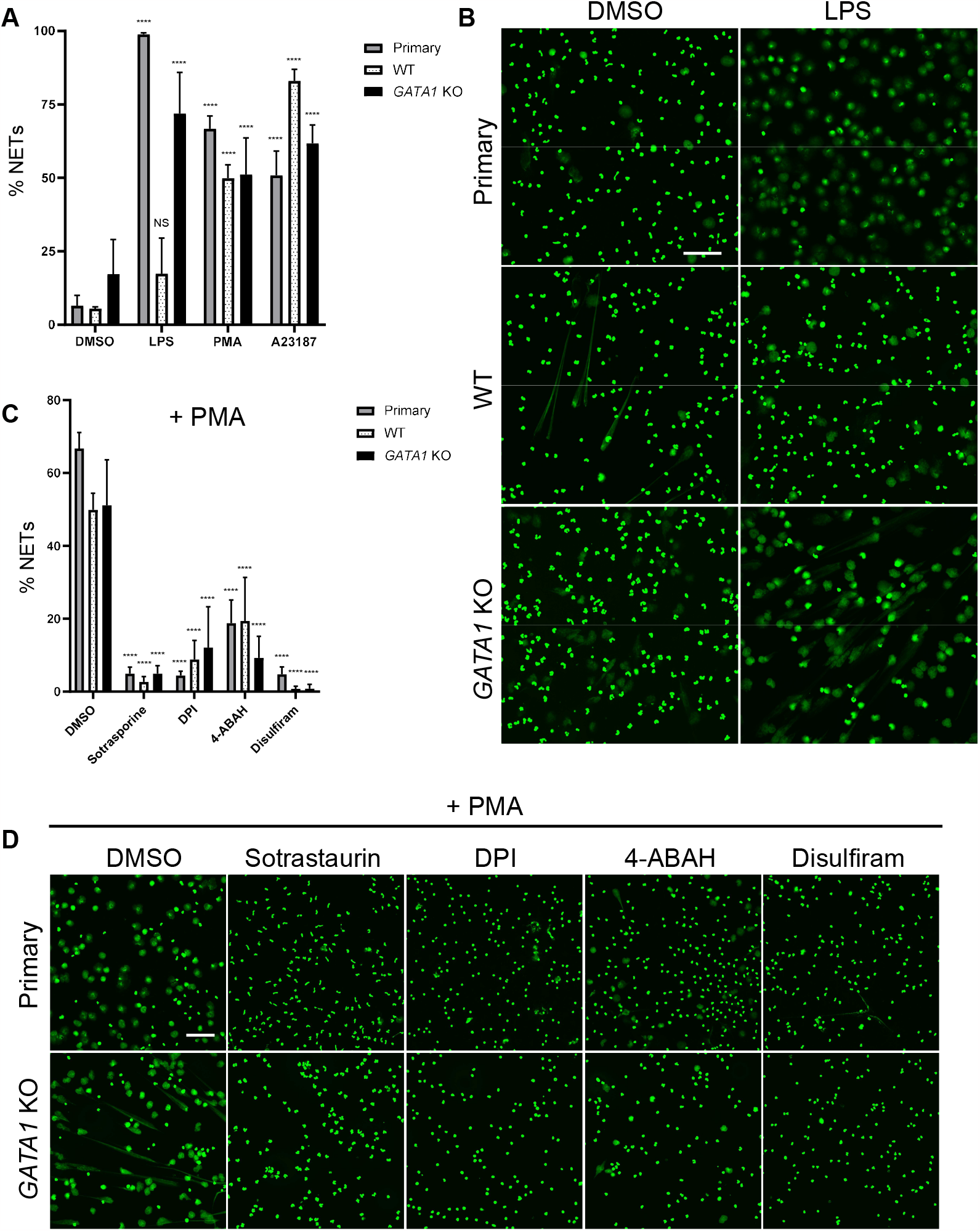
*GATA1* KO iNeutrophils form NETs like primary neutrophils. (A) Percentage of primary neutrophils, WT and *GATA1* KO iNeutrophils undergoing NET formation after 3 hours treatment with DMSO, 25 μg/ml LPS, 50 nM PMA or 5 μM A23187 on Day 19 (n = at least 3 independent experiments and 3 independent donors; mean ± SD). ****P < 0.0001 as compared to unstimulated DMSO controls determined by two-way ANOVA. (B) Fluorescent confocal microscopy of primary neutrophils’, WT and *GATA1* KO iNeutrophils’ nuclei (green) after treatment with DMSO or 25 μg/ml LPS for 3 hours on Day 19. Scale bar = 100 μm. (C) Percentage of primary neutrophils, WT and *GATA1* KO iNeutrophils undergoing NET formation after 1 hour pre-incubation with DMSO, 20 μM sotrastaurin, 20 μM DPI, 100 μM 4-ABAH or 20 μM disulfiram followed by 3 hours of stimulation with 50 nM PMA on Day 19 (n = at least 3 independent experiments and 3 independent donors; mean ± SD). ****P < 0.0001 as compared to PMA stimulated cells without inhibition determined by two-way ANOVA. (D) Fluorescent confocal microscopy of primary neutrophils’ and GATA1 KO iNeutrophils’ nuclei (green) after 1 hour pre-incubation with DMSO, 20 μM sotrastaurin, 20 μM DPI, 100 μM 4-ABAH or 20 μM disulfiram followed by 3 hours of stimulation with 50 nM PMA on Day 19. Scale bar = 100 μm.

While LPS is a well-described, physiologically relevant NET stimulant, it failed to induce significant NET formation in WT iNeutrophils (17 ± 12%) relative to DMSO controls (6 ± 1%), highlighting a severe limitation with this cell model (Figures 5A and 5B). Importantly, we observed that *GATA1* KO addresses this gap through restoring sensitization to LPS and generating significant NETs (72 ± 14%) relative to DMSO controls (17 ± 12%).

### NET formation can be inhibited in *GATA1* KO iNeutrophils using small molecules as in primary neutrophils

While PMA is non-physiologic, it is a commonly used tool to study NETs *in vitro* because it reliably activates relevant pathways (39). Like many physiologic NET forming stimulants, PMA activates PKC and subsequently generates ROS through the NADPH oxidase (NOX) complex. This releases MPO which helps decondense chromatin and expel DNA into the extracellular environment through the pore-forming protein Gasdermin D (40–42). We investigated the fidelity of this pathway in the WT and *GATA1* KO iNeutrophils using the PKC inhibitor sotrastaurin, the NOX inhibitor diphenylene iodonium (DPI), the MPO inhibitor 4-aminobenzoic acid hydrazide (4-ABAH) and the proposed Gasdermin D inhibitor, disulfiram. In line with primary neutrophils, NET formation in both the WT and *GATA1* KO iNeutrophils was significantly reduced after pre-treatment with these selective inhibitors (Figures 5C, 5D and S4B). These results demonstrate that the *GATA1* KO iNeutrophils respond to known NET inhibitors and can be used in screens to find novel small-molecule NET inhibitors.

### *GATA1* KO iNeutrophils can be genetically edited and used for target validation

*CYBB* encodes p91^phox^, a component of the multi-protein NADPH complex that is critical for NOX-dependent NET formation (39,43). To test whether *GATA1* KO iNeutrophils could be leveraged for functional genomic approaches, we knocked out *CYBB* using CRISPR/Cas9 in the *GATA1* KO hESCs, differentiated the cells to Day 19 iNeutrophils and stimulated the cells with PMA to induce NETs. Upon stimulation with PMA, 44 ± 13% of *GATA1* KO iNeutrophils with intact p91^phox^ (control gRNA) generated NETs compared to 6 ± 4% without p91^phox^ (Figures 6A and 6B). These results establish that our *GATA1* KO iNeutrophils form NETs with diverse stimuli like primary neutrophils, and that NET formation can be inhibited pharmacologically and genetically. We conclude that *GATA1* KO iNeutrophils overcome a major limitation associated with primary neutrophils, by enabling the identification and validation of targets modulating neutrophil functions.

**Fig. 6.**
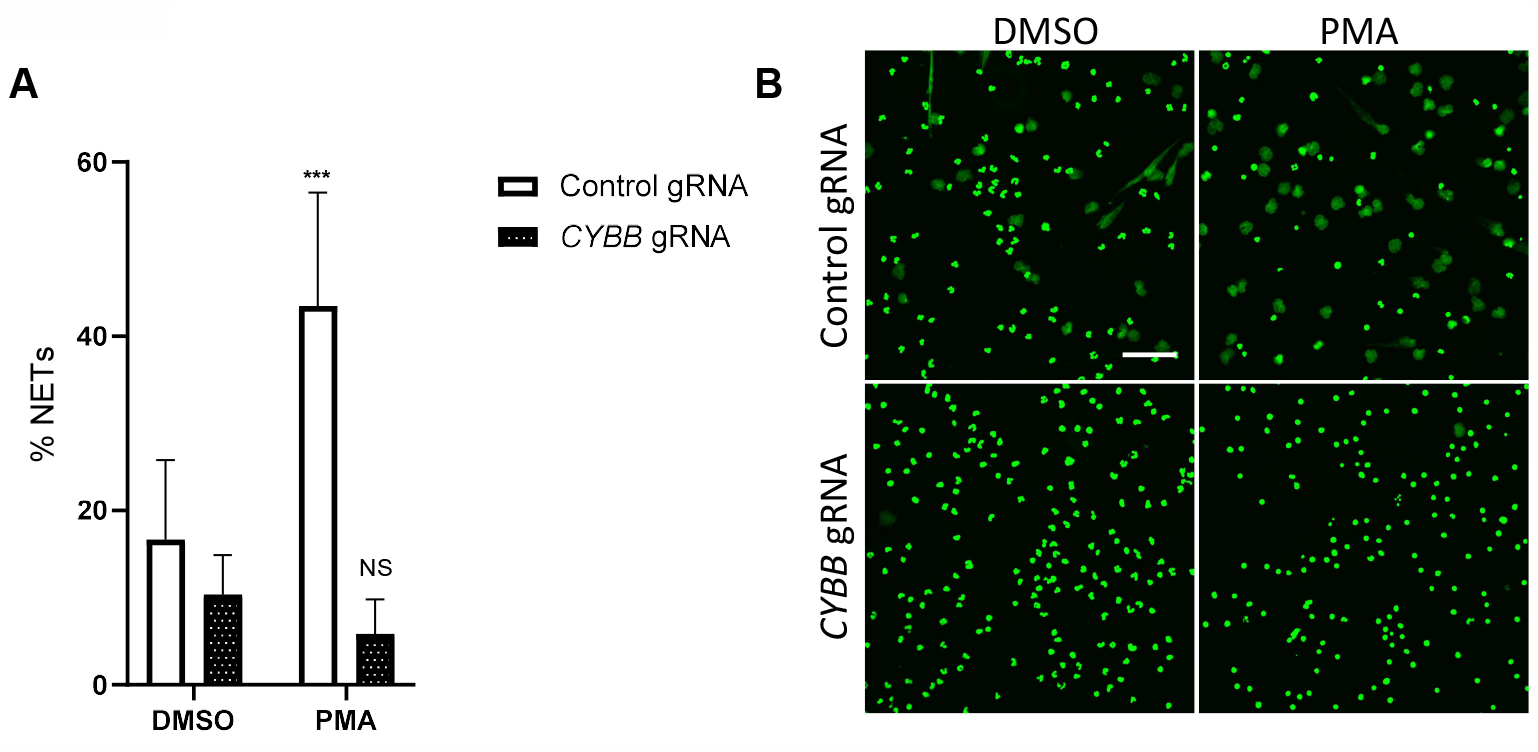
NET target validation in *GATA1* KO iNeutrophils. (A) Percentage of non-targeting (control gRNA) and *CYBB* gRNA lentiviral transduced *GATA1* KO iNeutrophils undergoing NET formation after 3 hours of treatment with DMSO or 50 nM PMA on Day 19 (n = at least 4 independent experiments; mean ± SD). ***P < 0.001 as compared to unstimulated DMSO controls determined by two-way ANOVA (B) Fluorescent confocal microscopy of control gRNA and *CYBB* gRNA transduced *GATA1* KO iNeutrophils’ nuclei (green) after stimulation with DMSO or 50 nM PMA for 3 hours. Scale bar = 100 μm.

## Discussion

In this study, we developed a novel method that dramatically increases the efficiency of differentiation, maturity, and functionality of hPSC-derived Neutrophils. Our method was adapted from previously published protocols to generate hematopoietic progenitors, with the addition of G-CSF between Day 12 and 18 to establish a granulocyte program, generating WT iNeutrophils. Recent reports optimizing the generation of iNeutrophils utilize gene overexpression to overcome differentiation challenges and deficiencies in host defense functions (37, 38). Overexpression and modification of genes can enhance iNeutrophil behavior *in vitro*, but strays from primary neutrophils in ways that may not be readily apparent. These methods also require genetic manipulation during each round of cell production, adding delivery challenges and/or FACS to purify targeted cells. Unlike these protocols, our method is amenable to engineering at the self-renewing hPSC stage where modified cells can be expanded and banked for further use.

Flow cytometry analysis of WT iNeutrophils confirmed previous observations that they are composed of two distinct populations: one characterized by an immunophenotype typical of primary neutrophils, and the other of either non-neutrophil granulocytes or hematopoietic progenitors. Sorting these two populations and comparing gene expression of five granulocyte regulators revealed that the neutrophil-like population expressed low *GATA1* while the non-neutrophil population highly expressed *GATA1*.

From this observation, we hypothesized that knocking out *GATA1* in the hESCs before differentiation would push them towards neutrophils and away from other fates. hESCs with CRISPR/Cas9 deletion of *GATA1* expressed high levels of pluripotency genes, which were lost upon differentiation, consistent with the behavior and expression changes seen in WT cells. After hematopoietic induction, levels of the hematopoietic progenitor markers *SPI1* and *GFI1* rose in the Day 6 monolayer cells. Large numbers of cells began shedding off the supporting monolayer between Day 7 and Day 8, and floating *GATA1* KO cells on Day 12 expressed higher levels of neutrophil specific genes relative to the WT control, supporting our hypothesis that *GATA1* removal encourages a neutrophil program.

In line with these findings, flow cytometry analysis showed a very high degree of similarity between the *GATA1* KO iNeutrophils and primary neutrophils. Other methods report producing populations with roughly 50% CD11b positive cells, and low levels of CD66b (37). *GATA1* KO enhances neutrophil specification and produces greater than 95% CD11b+ and 85% CD66b+ cells, with 81% expressing all the neutrophil surface proteins tested, precluding time-consuming and resource-intensive sorting. Our method therefore constitutes a substantial improvement over previously described approaches.

Prior to this study, the formation of NETs in iNeutrophils has mostly been assessed using PMA, and while PMA does robustly activate specific NET pathways, it is not a physiologic stimulant. Additionally, PMA-stimulated ETs are not unique to neutrophils and occur in eosinophils (37,38). This suggests that PMA-stimulated ETs observed in cells made following previous iNeutrophil protocols (which generate heterogenous granulocyte populations) could be from non-neutrophil cells. Conversely, LPS is a physiologic bacterial cell wall component known to stimulate NETs *in vitro* in primary neutrophils, and by itself does not evoke DNA release in eosinophils (44). Furthermore, studies demonstrate differential production of ETs from neutrophils and eosinophils in human disease, stressing the mechanistic differences between the two cell types (45). Current iNeutrophil protocols generate heterogeneous populations of different granulocytes, and while these cells form PMA stimulated ETs, they likely do not capture the disease-relevant nuances of neutrophil-specific NETs. The relevance of these cells in NET studies is therefore limited. The *GATA1* KO iNeutrophil model overcomes these limitations by generating pure cultures of neutrophil-like cells that respond to diverse NET stimulants like their primary counterpart. This is highlighted by the restoration of NET formation after stimulation with LPS.

While screens using primary neutrophils have uncovered drugs that inhibit NET formation (41), the targets of these drugs remain extremely challenging to pinpoint. Even if a ligand partner is discovered, this does not rule out off-target modalities. For instance, a group using the potent neutrophil elastase inhibitor GW311616A concluded neutrophil elastase is critical for NET formation (46). Follow up work employing selective neutrophil elastase inhibitors and knock-out mice dispute these findings, suggesting GW311616A’s NET inhibition mechanism is likely off-target (47,48). Furthermore, validating targets using CRISPR/Cas9 knockouts in primary human cells is challenging due to the neutrophils’ extremely short lifespan *ex vivo*, and the use of classic gene silencing techniques such as siRNA or shRNA may not be effective in neutrophils, which are rather stable and transcriptionally silent. Through the deletion of *CYBB*, we show how our iNeutrophils can provide a clean method to quickly validate targets without the uncertainty of compound off-target effects.

While targeting overactive NETs in disease is therapeutically attractive, interfering with other host-defense activities like phagocytosis and ROS release leave patients vulnerable to infections (49,50). Because *GATA1* KO iNeutrophils retain their other host defense capabilities, NET target knockout cells can serve as a tool to address the impact on these critical functions.

In conclusion, our differentiation method overcomes the limitations of previously published protocols by generating at-scale numbers of cells that are similar to primary neutrophils in form and function. COVID-19 highlights the link between dysregulated neutrophils and disease and stresses the therapeutic benefit of inhibiting unbalanced NETs. Our improved method accurately models NETs and enables large-scale screens and target validation in ways that were impossible using previous differentiation methods or primary neutrophils.

## Acknowledgements

Matthias Mueller, Rob Maher, Haoyuan Liu, Eric Shikatani, Gianfranco de Pascale, Katie Worringer

