## Supplemental Figure 1 for "*GATA1* knockout in human pluripotent stem cells generates enhanced neutrophils to investigate extracellular trap formation"

```
GATA1 KO   TCTGTCCTCGCAGGTTAATCCCCAGAGGCTCCATGGAGTTCCCTGGCCTGGGGTCCCTGG
           ||||||||||||||||||||||||||||||||||||||||||||||||||||||||||||
Reference  TCTGTCCTCGCAGGTTAATCCCCAGAGGCTCCATGGAGTTCCCTGGCCTGGGGTCCCTGG

GATA1 KO   GGACCTCAGAGCCCCTCCCCCAGTTTGTGGA-------------TCCTCCACACCAGAAT
           |||||||||||||||||||||||||||||||             ||||||||||||||||
Reference  GGACCTCAGAGCCCCTCCCCCAGTTTGTGGATCCTGCTCTGGTGTCCTCCACACCAGAAT

GATA1 KO   CAGGGGTTTTCTTCCCCTCTGGGCCTGAGGGCTTGGATGCAGCAGCTTCCTCCACTGCCC
           ||||||||||||||||||||||||||||||||||||||||||||||||||||||||||||
Reference  CAGGGGTTTTCTTCCCCTCTGGGCCTGAGGGCTTGGATGCAGCAGCTTCCTCCACTGCCC
```

NCBI BLAST alignment screen shot of *GATA1* KO hESC vs reference (WT) DNA. A 13 base-pair deletion in *GATA1* exon 1 generates a frame shift mutation.
