## Supplemental Figure 2 for "*GATA1* knockout in human pluripotent stem cells generates enhanced neutrophils to investigate extracellular trap formation"

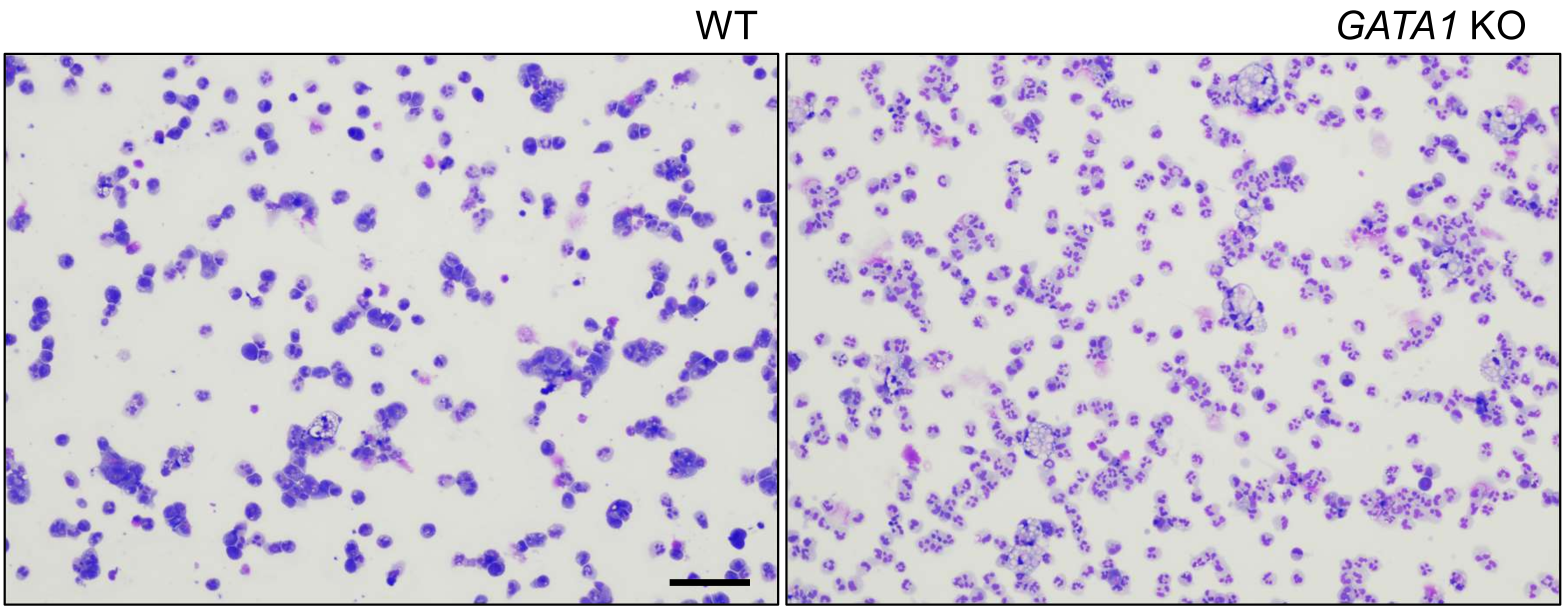


Hematoxylin and eosin images of WT and *GATA1* KO iNeutrophils harvested on Day 19. Scale bar = 100 μm.
