## Supplemental Figure 3 for "*GATA1* knockout in human pluripotent stem cells generates enhanced neutrophils to investigate extracellular trap formation"

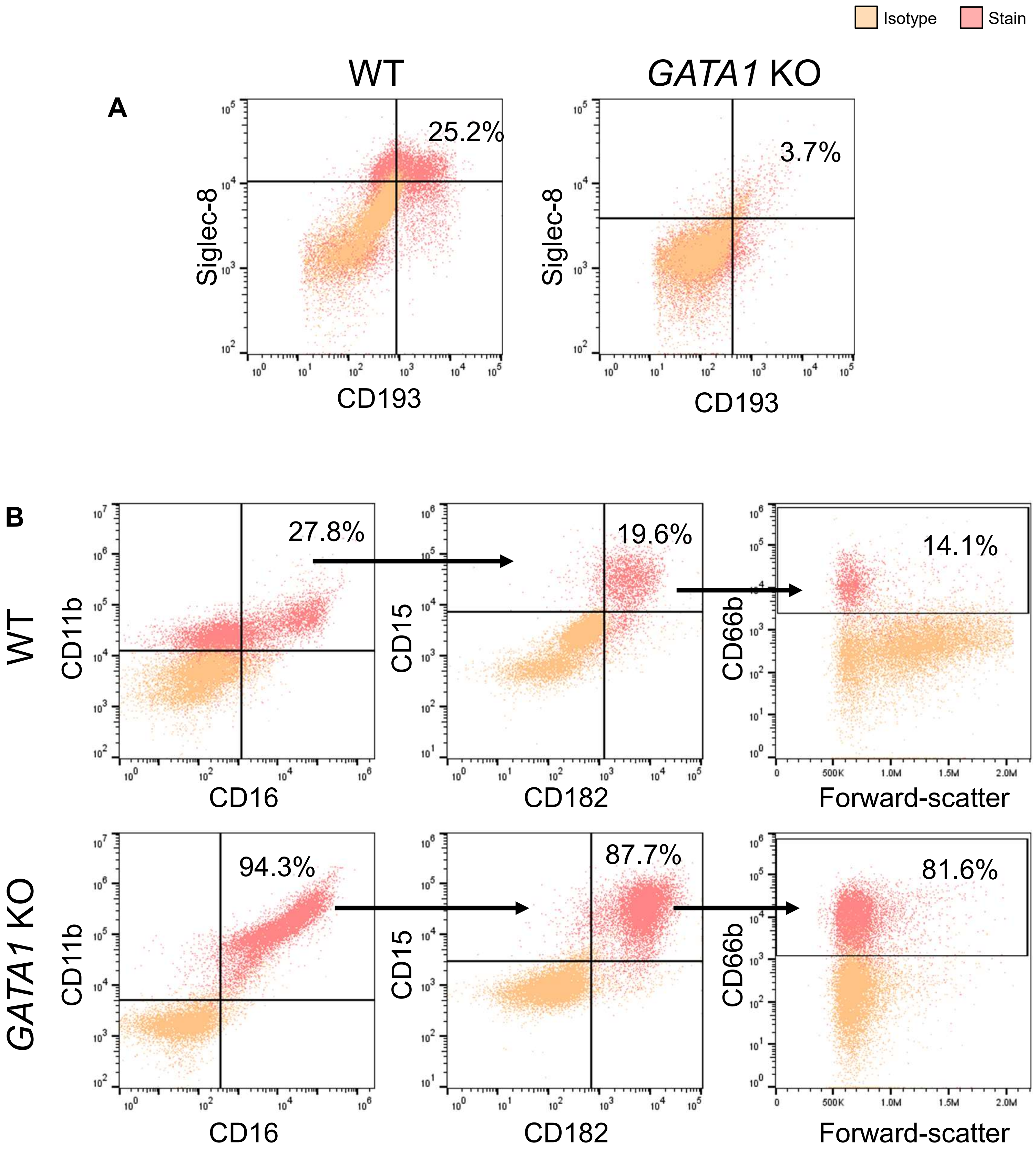


(A) Flow cytometry analysis of WT and *GATA1* KO iNeutrophils co-stained using antibodies against non-neutrophil granulocyte surface markers CD193 and Siglec-8 (B) and antibodies against neutrophil markers CD11b, CD16, CD15, CD182 and CD66b on Day 19. Plots show isotype control (orange) and corresponding specific antibody (red) scatter plots. Percentages represent total cells harvested, not just cells from previous gate.
