## Supplemental Figure 4 for "*GATA1* knockout in human pluripotent stem cells generates enhanced neutrophils to investigate extracellular trap formation"

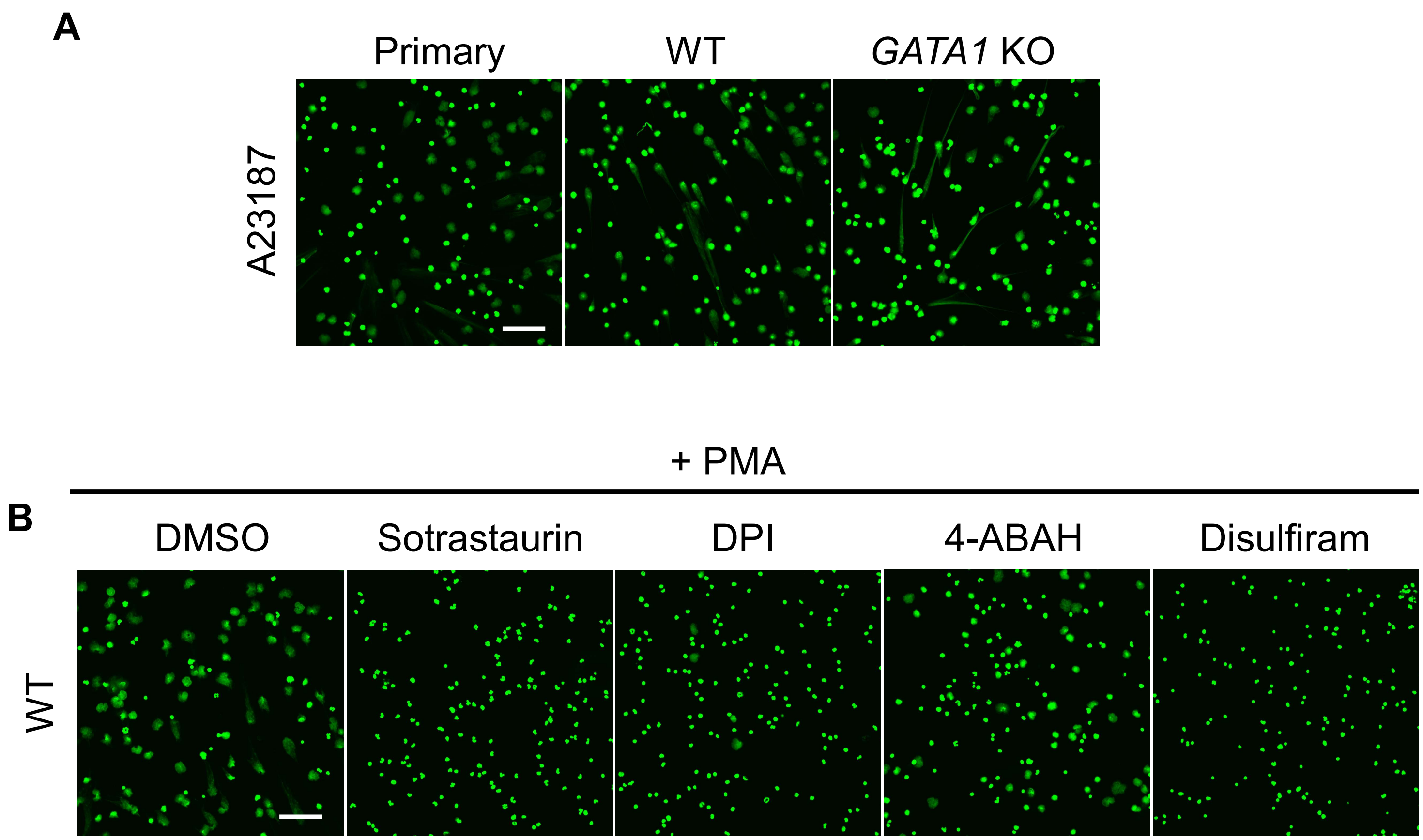


(A) Fluorescent confocal microscopy of primary neutrophils, WT and *GATA1* KO iNeutrophils' nuclei (green) after treatment with 5 μM A23187 for 3 hours on Day 19. (B) Fluorescent confocal microscopy of WT iNeutrophils' nuclei (green) after 1 hour pre-incubation with DMSO, 20 μM sotrastaurin, 20 μM DPI, 100 μM 4-ABAH or 20 μM disulfiram followed by 3 hours of stimulation with 50 nM PMA on Day 19. Scale bar = 100 μm.
