## Supplemental Table 1 for "*GATA1* knockout in human pluripotent stem cells generates enhanced neutrophils to investigate extracellular trap formation"

**A**

| Antigen | Fluorophore | Manufacturer | Host | Isotype | Catalog # |
|---|---|---|---|---|---|
| CD33 | APC | BioLegend | Mouse | IgG1 | 366605 |
| CD45 | FITCI | BioLegend | Mouse | IgG1 | 368508 |
| CD15 | BV510 | BD Biosciences | Mouse | IgG1 | 563141 |
| CD66b | PECy7 | eBioscience | Mouse | IgM | 25-0666-42 |
| CD11b | APC | BD Biosciences | Rat | IgG2b | 561690 |
| CD16 | APCCy7 | BD Biosciences | Mouse | IgG1 | 557758 |
| CD182 | PE | BioLegend | Mouse | IgG1 | 320705 |
| CD193 | PE | BioLegend | Mouse | IgG2b | 310705 |
| CD49d | FITCI | BioLegend | Mouse | IgG1 | 304315 |
| Siglec 8 | APC | BioLegend | Mouse | IgG1 | 347105 |

**B**

| Isotype control | Fluorophore | Manufacturer | Catalog # |
|---|---|---|---|
| IgG1 | APC | BioLegend | 400122 |
| IgG1 | FITCI | BioLegend | 400108 |
| IgG1 | BV510 | BD Biosciences | 562946 |
| IgM | PECy7 | eBioscience | 25-4752-81 |
| IgG2b | APC | BD Biosciences | 553991 |
| IgG1 | APCCy7 | BD Biosciences | 557873 |
| IgG1 | PE | BioLegend | 400113 |
| IgG2b | PE | BioLegend | 401207 |

(A) Flow cytometry specific antibodies and (B) isotype controls.
