## Supplemental Table 2 for "*GATA1* knockout in human pluripotent stem cells generates enhanced neutrophils to investigate extracellular trap formation"

| Gene | Catalog # | Marker |
|---|---|---|
| *SPI1* | hs02786711_m1 | Hematopoietic transcription factor |
| *GFI1* | hs00382207_m1 | Hematopoietic transctiption repressor |
| *AZU1* | hs00156049_m1 | Neutrophil granular protein |
| *AQP9* | hs01033361_m1 | Neutrophil water / glycerol channel protein |
| *ELANE* | hs00236952_m1 | Neutrophil granular protein |
| *MPO* | hs00165162_m1 | Neutrophil granular protein |
| *CLC* | hs00171342_m1 | Eosinophil / basophil lysophospholipid |
| *OCT4* | hs00999632_g1 | Pluripotency transcription factor |
| *Nanog* | hs02387400_g1 | Pluripotency transcription factor |
| *CEBPD* | hs00270931_s1 | Hematopoietic transcription factor |
| *CEBPE* | hs00357657_m1 | Hematopoietic transcription factor |
| *GATA1* | hs01085823_m1 | Hematopoietic transcription factor |
| *GAPDH* | hs99999905_m1 | Housekeeping gene |

TaqMan probes used in RT-qPCR
