## Supplemental Table 3 for "*GATA1* knockout in human pluripotent stem cells generates enhanced neutrophils to investigate extracellular trap formation"

| Marker | Cell type | Primary % + | WT % + | *GATA1* KO % + |
|---|---|---|---|---|
| CD182 | Granulocyte | 98 | 83 | 94 |
| CD11b | Granulocyte | 100 | 88 | 98 |
| CD15 | Neutrophil | 99 | 50 | 95 |
| CD16 | Neutrophil | 100 | 29 | 95 |
| CD66b | Granulocyte | 98 | 22 | 88 |
| Siglec-8 | Eosinophil | 4 | 51 | 5 |
| CD193 | Eosinophil | 6 | 37 | 8 |
| CD49d | Non-neutrophil granulocyte | 3 | 26 | 6 |

Cell surface markers and expression percentages
